# Molecular Basis of Sodium Channel Inactivation

**DOI:** 10.1101/2025.05.22.655422

**Authors:** Yichen Liu, Jason D. Galpin, Christopher A. Ahern, Francisco Bezanilla

**Author notes:** Corresponding author: Francisco Bezanilla.

## Abstract

Voltage-gated sodium channels initiate action potentials and control electrical signaling throughout the animal kingdom. Fast inactivation is an essential auto-inhibitory mechanism and requisite component of sodium channel physiology. Recent structural and electrophysiological results are inconsistent with the canonical “ball and chain” model of fast inactivation thus necessitating an updated theoretical framework. Here, we use encoded fluorescence spectroscopy and high-resolution electrophysiology to capture key steps in the fast inactivation mechanism, from voltage-sensor activation to pore occlusion, an ultra-fast process which occurs in less than 2 milliseconds. Upon depolarization, activation of the domain IV voltage sensor initiates cytoplasmic DIII_DIV linker movement and quickly repositions the IFM motif into a hydrophobic pocket adjacent to the pore. This triggers a structural rearrangement of the pocket. The phenylalanine of the IFM motif contacts the pore-forming helices via a hydrophobic interaction with S6 of DIV and an aromatic/hydrophobic interaction with S6 of DIIII. These two interactions occur only after both S6 segments rotate, thus exposing the hydrophobic gate into the pore producing the fast inactivation. Based on the current results, we propose an alternative “lock and key” model to explain the molecular mechanism of fast inactivation.

## Introduction

The action potential is the elementary unit of bio-excitability. At the molecular level, voltage-gated sodium (Nav) channels are responsible for the rapid upstroke of the action potential. These channels are gated to an open, sodium conducting conformation by positive changes in transmembrane potential ^1–3^. During the rising phase of the action potential, Nav channels operate in a classical positive-feedback manner: opened by membrane depolarization, the Nav channels allow further Na^+^ entry with consequent membrane depolarization, thus resulting in a cascading, all-or-nothing action potential. In this positive feedback loop, fast inactivation is an essential built-in auto-inhibitory mechanism to terminate the cascade and serves as a molecular “kill switch”. Milliseconds after the channels are open during activation, fast inactivation drives the channels into the nonconductive, fast inactivated state, facilitating the repolarization by voltage gated potassium channels. Acquired or inherited abnormalities in fast inactivation can result in severe pathological states ^4^. Additionally, due to the prevalence and diversity of Nav channels, these pathological states can manifest in numerous tissue types with a wide range of symptoms ^5–9^. Hence, a molecular understanding of the fast inactivation mechanism is of fundamental importance.

Originally proposed almost 50 years ago, the “ball and chain” model predicts that an intracellular located “inactivation particle” would bind to the open channel at the pore and physically block the ionic conduction ^10^. Later, the identification of the amino acid triad, Isoleucine-<u>Ph</u>enylalanine-<u>M</u>ethionine (the IFM motif) in the cytoplasmic domain III and domain IV (DIII_DIV) linker as the inactivation particle materialized the abstract concept and provided a molecular framework for fast inactivation ^11^. Numerous studies have since provided strong evidence for this model and consequently, the “ball and chain” model has been largely accepted as the textbook explanation for fast inactivation in Nav channels^12–16^.

However, upon the advent of the cryogenic electron microscopy mediated “resolution revolution” in structural biology, discrepancies emerged between the old models and new experimental results ^17^. As of today, essentially all of the available eukaryotic Nav channel structures, likely in the inactivated state due to the lack of external electric field, consistently demonstrate that the IFM motif, the inactivation particle, does not block the permeation pathway as expected ^18–21^. Instead, the motif docks in an adjacent hydrophobic pocket between the pore and voltage-sensor domains, a stark contradiction to the canonical pore-blocking “ball and chain” model. Our recent work began to address this inconsistency and identified the inactivation gate, a pore-lining double-layered hydrophobic ring at the intracellular mouth of the pore that blocks Na^+^ ion permeation in the inactivated state, instead of the IFM motif ^22^. However, as this is only the final step, additional modifications are necessary for the “ball and chain” model to accommodate the current structural and physiological results. The inactivating particle (IFM motif) is necessary for inactivation but the mechanistic linkage between voltage-sensor activation, the docked IFM motif and the newly described gate remains poorly resolved. Thus, the lack of molecular description and theoretical framework of the fast inactivation process once again represents a major conceptual gap in Nav channel biophysics and physiology.

Here, we present the full sequence of conformational changes during the fast inactivation process in mammalian Nav channels. The dynamic nature of the molecular rearrangements between the IFM binding and the gate closing requires techniques that can detect these changes while observing the actual function of the channel. To this end we have used site-directed fluorescence in voltage clamped conditions. With our improved incorporation protocol and optical recording setup, we introduced an environmentally sensitive fluorescent unnatural amino acid, ANAP, into various regions of the Nav channel and monitored the local molecular motions during fast inactivation by rapid fluorescence measurements. ANAP fluorescence recordings reported two types of movements at the IFM location during fast inactivation. A fast movement, triggered by the activation of voltage sensing domain (VSD) in domain IV (DIV), reflects the movement of the DIII_DIV linker region and posits the IFM motif into the appropriate location, conceptually like putting a key into a keyhole, that was exposed by the movement VSD of DIV. Coupled by hydrogen bonds between DIII_DIV linker and DIV S4_S5 linker, this fast component follows faithfully the movements of DIV VSD, via simultaneously measured gating currents, and proceeds prior to the closure of the fast inactivation gate. The other slower conformational change at the IFM motif triggers the fast inactivation. The aromatic side chain of the phenylalanine residue in the IFM motif forms interactions with two residues in DIII S6 and DIV S6, through T-shaped π-π and hydrophobic interactions. In tandem, these interactions propagate the conformational change from the linker to the pore and lead to the closure of the double-layered fast inactivation gate, similar to the movement of the bolt of the lock. Contrary to our expectations, modification to the interaction between the IFM motif and the hydrophobic pocket does not impede the binding of the motif itself, rather it appears to influence the effectiveness of the IFM motif at triggering fast inactivation, like creating mismatches between the key and the lock. With these current observations, we propose a “lock and key” model to describe the sequential conformational changes during fast inactivation in mammalian sodium channels.

## Results

### IFM Motif Undergoes Two Distinct Movements During Fast Inactivation

The IFM motif is evolutionarily conserved in Nav channels across the animal kingdom (Fig. 1A). Its importance for fast inactivation has been demonstrated repeatedly in the literature^14,23^. However, recent structural results have cast uncertainty as to the functional role IFM motif plays in the mechanism of fast inactivation. To describe the conformational changes at the IFM position and to understand its role in fast inactivation, we replaced the phenylalanine in the IFM motif with a fluorescent, environmentally sensitive unnatural amino acid, ANAP (F1304ANAP, numbering based on rat Nav 1.4)^24,25^. ANAP (3-(6-<u>a</u>cetyl<u>n</u>aphthalen-2-ylamino)-2-<u>a</u>mino<u>p</u>ropanoic acid) has a comparable size as tryptophan which allows minimum disturbance upon incorporation (Fig. 1B). More importantly, the emission spectrum of ANAP is modulated by the hydrophobicity of its immediate environment, making ANAP an ideal reporter of molecular movement ^25^ (Fig. 1B). Robust currents were seen from F1304ANAP, and no discernable current was seen in cells where no ANAP was supplied, thus demonstrating high fidelity of encoding (Fig. 1C). F1304ANAP channels showed minor slowing effects on fast inactivation, similar to tryptophan replacement^14^, and minimal impact in activation (Fig. 1D-F). Utilizing site-directed fluorimetry with cut-open voltage clamp technique, ionic current and fluorescence signal were resolved simultaneously^26,27^.

**Figure 1:**
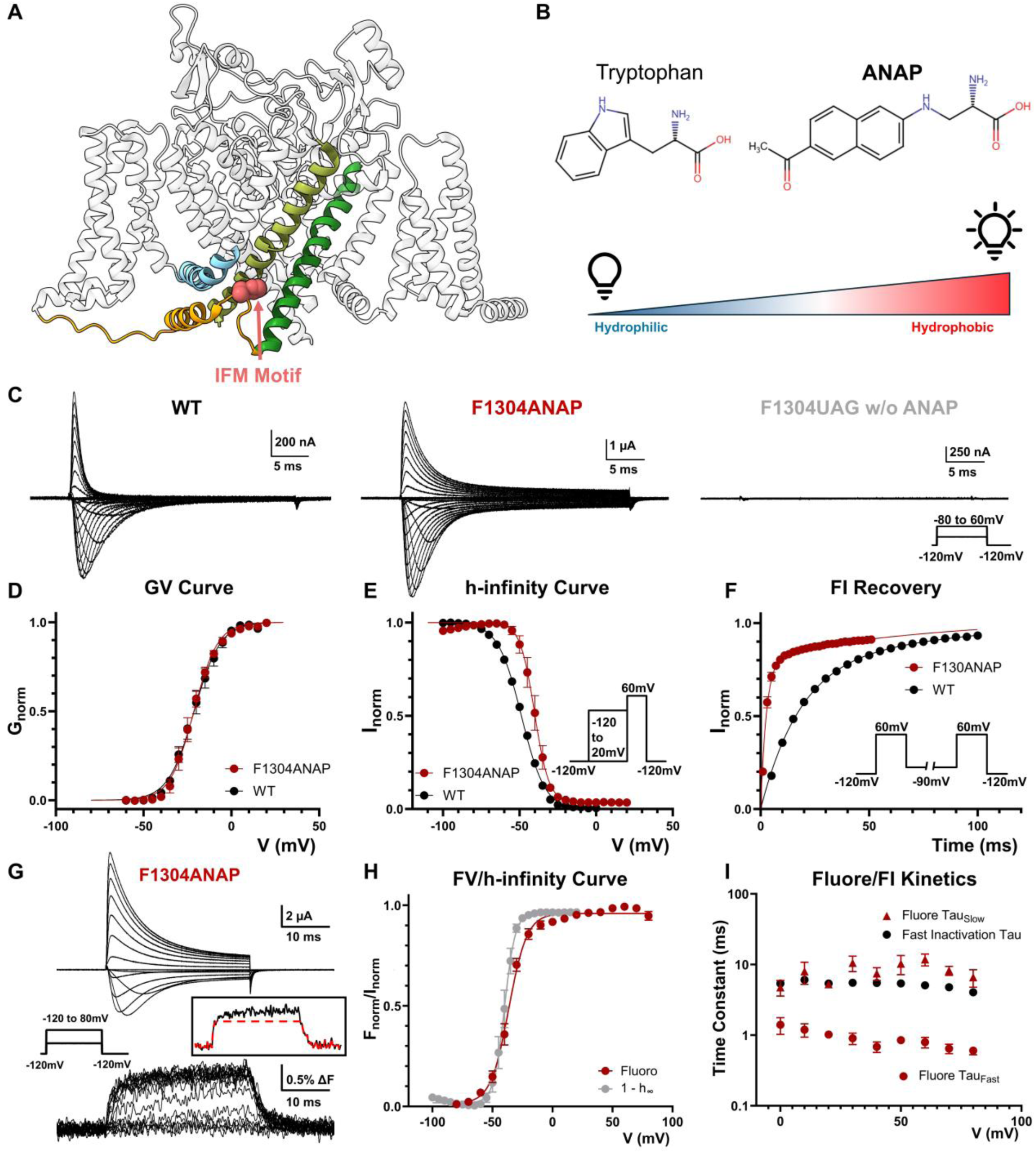
Incorporation of ANAP and IFM Movement during fast inactivation. **A)** Side view of putative inactivated Nav channel structure (PDB: 7XVF). IFM motif (in pink) resides in the DIII_DIV linker (yellow) and docks into a pocket away from the pore. **B)** Size of ANAP is comparable to tryptophan. With the filter set used in the study (details in Methods), the fluorophore is brighter in hydrophobic environment and dimmer in hydrophilic environment. **C)** Exemplar ionic traces of wild type (WT), F1304ANAP and F1304UAG when no ANAP was supplied. Incorporation of ANAP into IFM motif modestly slowed fast inactivation kinetics. Inset shows the voltage protocol. **D)** GV curve of WT and F1304ANAP. No significant difference was observed. N = 4 for WT and N = 5 for F1304ANAP. **E)** h-infinity curve for WT and F1304ANAP, with the latter showing a slightly right-shifted ΔV_half_ = ∼8.5mV. Inset shows the voltage protocol. N = 4 for WT and N = 5 for F1304ANAP. **F)** Recovery from fast inactivation at -90mV. F1304ANAP recovered much faster than the WT, suggesting a less stable inactivated state. N = 4 for WT and N = 5 for F1304ANAP. Inset shows the voltage protocol. **G)** Exemplar family of traces of simultaneously recorded ionic current and fluorescence signal from F1304ANAP. An increase of fluorescence intensity indicates a transition from a relative hydrophilic environment to a more hydrophobic one. Two kinetic components can be distinguished from the fluorescence signal at IFM motif. Left inset shows the voltage protocol and right inset shows the fluorescence signal at 60mV fitted with only one fast component. **H)** Comparison of steady state fluorescence (FV) curve and h-infinity curve. H-infinity curve is shown as 1-(h-infinity) curve which closely tracks with the FV curve. N = 5. **I)** Comparison of fluorescence signal and fast inactivation kinetics. The fast fluorescence signal is significantly faster than the kinetics of fast inactivation. The slow component, on the other hand, follows closely the fast inactivation kinetics. N = 5 for fluorescence signal and N = 4 for ionic current. Data is shown as mean ± SEM.

Fluorescence signals increased upon depolarization, indicating an increase in the environmental hydrophobicity seen by the side chain of ANAP (Fig. 1G). This is consistent with the notion that during fast inactivation the IFM motif transitions from a hydrophilic to more hydrophobic environment. The overlap of the steady-state fluorescence (FV) curve and fast inactivation, tracked via 1-(h-infinity) curve (Fig. 1H), indicates that the movement of the IFM motif directly reports the fast inactivation state of the Nav channels. Additionally, the off-fluorescence signal closely tracks the recovery from inactivation (Supple. Fig. 1). Interestingly, the fluorescence signals that originate from F1304ANAP channels exhibit two kinetic components, suggesting two distinct conformational changes in the vicinity of the IFM motif occurred during fast inactivation (Fig. 1G, inset). Comparison between the fast inactivation kinetics and the fluorescence signal kinetics revealed that the fast time constant of fluorescence signal was significantly faster than the fast inactivation kinetics. The slow component on the other hand, closely followed the development of fast inactivation (Fig. 1I).

### Fast Movement at IFM Motif Reflects DIII_DIV Linker Movement Following DIV VSD Activation

To further investigate the nature of conformational dynamics at the IFM motif during fast inactivation, we monitored the movement of the IFM-bearing DIII_DIV linker by incorporating ANAP at L1319 position (L1319ANAP, Fig. 2A). ANAP incorporation at position 1319 was well tolerated (Supple. Fig. 2) and opposed to the F1304 position, L1391ANAP channels showed a marked decrease in the fluorescence intensity upon depolarization, suggesting a transition to more hydrophilic environment (Fig. 2B). This is consistent with the existing structural results since L1319 is depicted as being completely exposed in aqueous environment in the fast inactivated structure (Fig. 2A). More interestingly, there is only one kinetic component in the fluorescence signal at L1319 position. This component showed similar time constants as the fast movement seen in F1304ANAP (Fig. 2C, D). Like the fast fluorescence signal in F1304ANAP channels, the fluorescence signal recorded in L1319ANAP is faster than the development of fast inactivation itself (Fig. 2E). To better understand the origin of this signal, we simultaneously recorded fluorescent signals and gating currents in L1319ANAP channels (Fig. 2F). Upon depolarization, gating charges in voltage sensors rapidly traverse the electric field, creating a fast, transient current, the gating current ^28,29^. Given that gating currents track the movement of VSDs, this can be correlated to the observed fluorescence signals that originate from movements at the cytoplasmic III-IV linker. Gating current recordings from L1319ANAP channels revealed that the voltage dependent gating charge movement, shown by the QV curve, overlaps closely with the steady-state FV curve (Fig. 2G). Despite the relative distance between the VSD and III-IV linker, the movement seen at L1319 linker region is more closely related to the voltage sensor movements than to fast inactivation. More specifically, the DIV VSD movement appears to directly trigger the DIII_DIV linker movement. When we compared the kinetics of the gating current and the fluorescence recordings, we discovered that the fluorescence signal at position 1319 explicitly followed the slow component of the gating current (Fig. 2H). Due to the asymmetry in Nav channels, voltage sensors from different domains translocate their charges at different rates ^30,31^. Particularly, the activation of the DIV VSD is significantly slower than the other VSDs, giving rise to the slow component of the gating current in Nav channels^30,32^. The shared kinetics between fluorescence signal at position 1319 and the DIV VSD movement strongly suggests the activation of voltage sensor in DIV initiates the DIII_DIV linker movement during fast inactivation.

**Figure 2:**
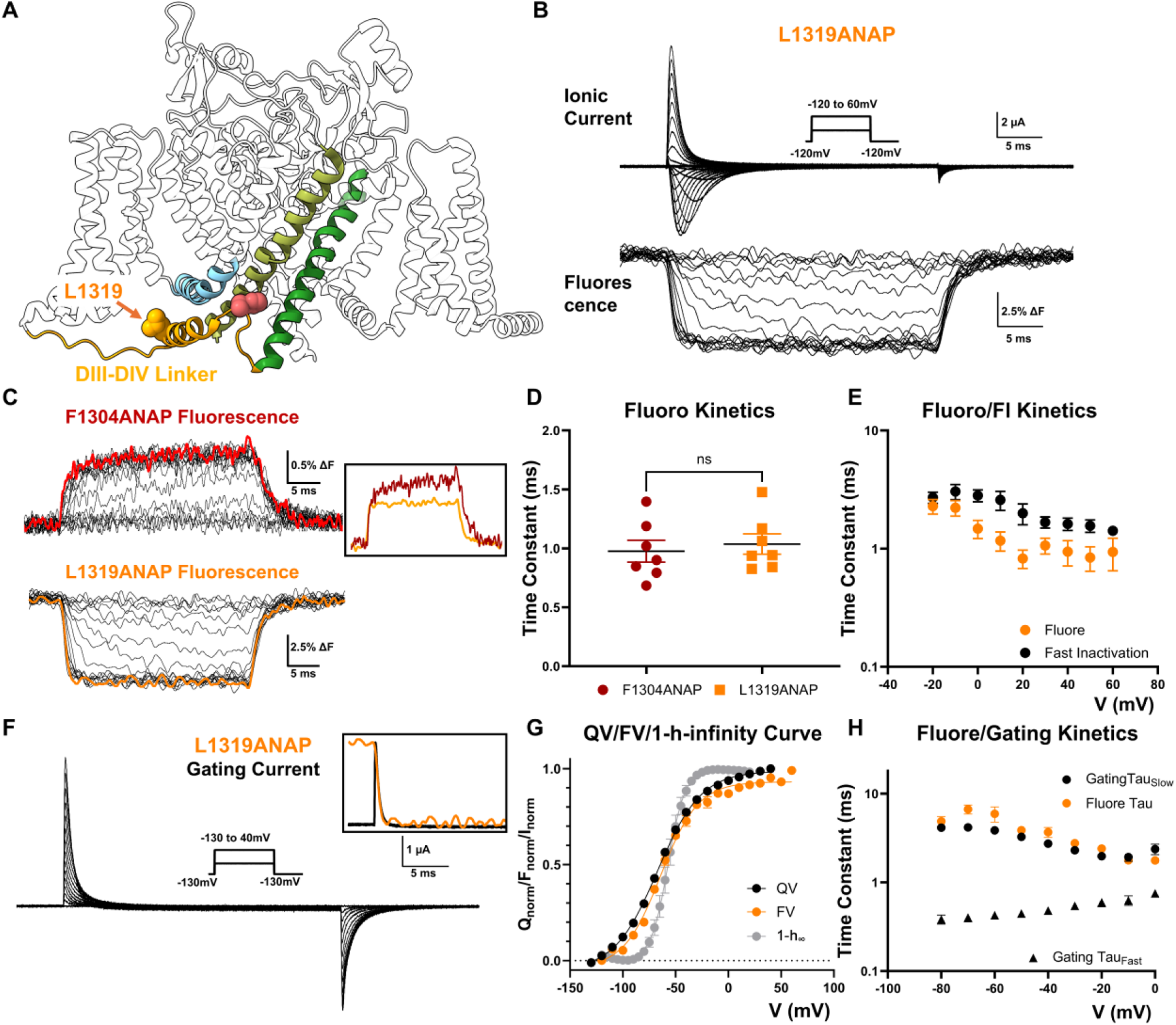
DIII_DIV linker movement gives rise to the fast component following DIV VSD movement. **A)** To monitor the movement of DIIII_DIV linker region during fast inactivation, ANAP was incorporated at L1319 position, in the short alpha helix. **B)** Exemplar traces of L1319ANAP ionic current and fluorescence signal. Inset shows the voltage protocol. **C)** Comparison of fluorescence signal from F1304ANAP and L1319ANAP. Inset shows a scaled comparison of the fluorescence signals at both positions at 40mV. **D)** The fast component of the fluorescence signal from F1304ANAP is not significantly different from the signal at position 1391 at 60mV. **E)** Comparison of the fluorescence signal and the fast inactivation kinetics of L1319ANAP. The fluorescence signal was always faster than the fast inactivation development. N = 7 for fluorescence signal and N = 5 for ionic current. **F)** Exemplar traces of gating current measured from L1391ANAP. Inset shows the voltage protocol. Framed inset shows a scaled comparison of fluorescence signal and gating current at 40mV where gating charge moved in a fast single exponential manner. **G)** Comparison of voltage dependent gating charge movement (QV curve), FV curve of L1319ANAP and (1-h)-infinity curve. Clearly, the fluorescence signal is following more closely the gating charge movement. N = 9 for QV, N = 10 for FV and N = 6 for h-infinity curve. **H)** Comparison of fluorescence kinetics of L1319ANAP and gating charge movement. The fluorescence kinetics is similar to the slow component of gating charge movement, contributed mostly by DIV VSD movement. N = 9. Data is shown as mean ± SEM.

### Backbone Hydrogen Bonding Couples the DIII_DIV Linker and DIV VSD Movement

Since the DIII_DIV linker movement closely correlates with the activation of DIV VSD, there must exist interactions that couple these two structural components. A close structural analysis of this region revealed the existence of two hydrogen bonds between the DIII_DIV linker and the DIV S4_S5 linker (Fig. 3A). Specifically, residue N1477 at the “elbow region” of the DIV S4_S5 linker forms a hydrogen bond directly with the F1304 in the bound IFM motif and reciprocally, Q1309 at the DIII_DIV linker forms the other hydrogen bond with P1473 in DIV (Fig. 3A). Interestingly, both hydrogen bonds are formed between the nitrogen in the primary amide in the side chain and the carbonyl oxygen in the backbone. Removal of the primary amide in the side chain in both positions result in impaired fast inactivation (Fig. 3B). In the case of Q1309L and N1477D, the channels exhibit an “IQM” type phenotype with most of the fast inactivation absent^11,31^ (Fig. 3C, D and F) without drastic change in GV curves (Fig. 3C). An analysis of the recovery from fast inactivation for Q1309A channels further demonstrates that the identified hydrogen bond is essential for the stabilization of the fast inactivated state (Fig. 3E). In the absence of this interaction, the channels recover from fast inactivation exceedingly fast. Evidently, these hydrogen bonds are important for fast inactivation and likely serve as the functional bridges between the DIV VSD and the intracellular linker. To further demonstrate this, we disrupted one of the hydrogen bonds and investigated its impact on DIII_DIV linker movement. In L1319ANAP_Q1309A, fast inactivation becomes slower and incomplete (Fig. 3I). The fluorescence signal became much smaller (Fig. 3I) and more importantly, progressed slower and recovered faster compared to L1319ANAP (Fig. 3F, G), demonstrating the identified interactions are indeed involved in coupling between the voltage sensor in DIV (via the DIV S4-S5 linker) and DIII_DIV linker.

**Figure 3:**
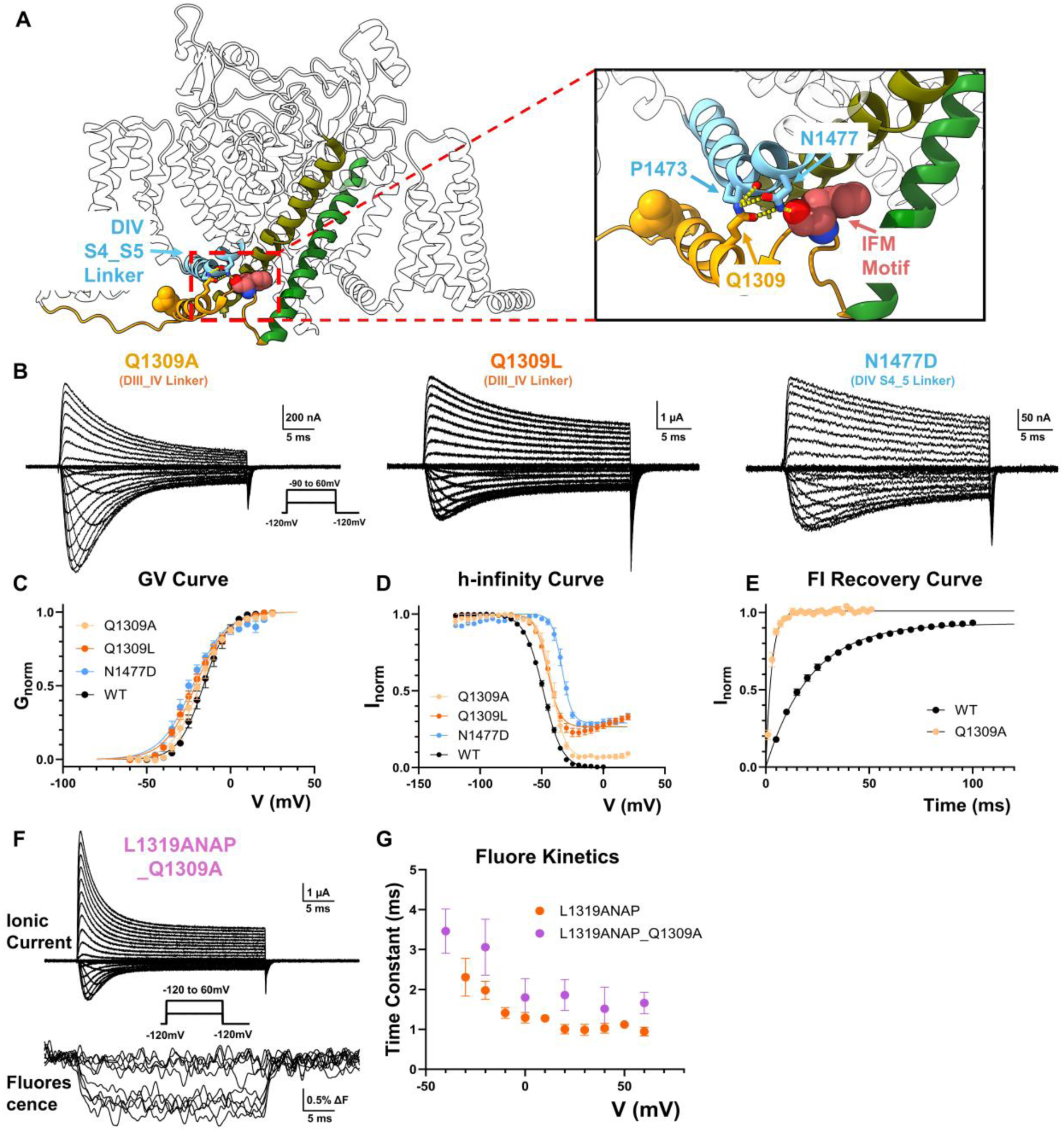
Backbone hydrogen bonds couple DIII_DIV linker movement to DIV VSD movement. **A)** Structural analysis reveals a network of hydrogen bonds between DIII_DIV linker (yellow) and DIV S4_S5 linker (cyan). Q1309 from the DIII_DIV linker forms a potential hydrogen bond with the backbone carbonyl group of P1473 DIV S4_S5 linker while N1477 in the same region forms directly a putative hydrogen bond with the backbone of the phenylalanine in IFM motif (pink). **B)** Exemplar ionic current from Q1309A, Q1309L and N1477D respectively. Inset shows the voltage protocol. **C)** GV curves for Q1309, Q1309L and N1477D. Activation of the mutant channels is not significantly altered compared to WT. N = 5 for Q1309A, N = 4 for Q1309L and N = 5 for N1477D. **D)** h-infinity curves for the indicated channels. N = 4 for Q1309A, N = 4 for Q1309L and N = 6 for N1477D. **E)** Q1309A recoveries from fast inactivation significantly faster than WT, suggesting a destabilized inactivated state. N = 3 for Q1309A. **F)** Example traces of ionic current and fluorescence signal from L1319ANAP_Q1309A. Inset shows the voltage protocol. **G)** Comparison of fluorescence kinetics of L1319ANAP and L1319ANAP_Q1309A. Mildly disrupting the hydrogen bond, L1319ANAP_Q1309A showed slowed fluorescence kinetics than L1319ANAP, suggesting the identified hydrogen bonds are important for the coupling of the DIV VSD and DIII_DIV linker. N = 6. Data is shown as mean ± SEM.

### IFM Motif Relays Its Conformational Changes to the Pore through Hydrophobic and T-Shaped π-π Interactions

With the molecular underpinning of the fast component elucidated, we set out to investigate the slower conformational change at the IFM motif that triggers fast inactivation. Our previous work demonstrated the specialized role of the pore-lining hydrophobic residues in DIII and DIV S6 in fast inactivation^22^ (Fig. 4A). Therefore, residues in DIII and DIV S6 that could form potential interactions with the IFM motif were screened through site-directed mutagenesis. The screening revealed two residues that were particularly important for fast inactivation.

**Figure 4:**
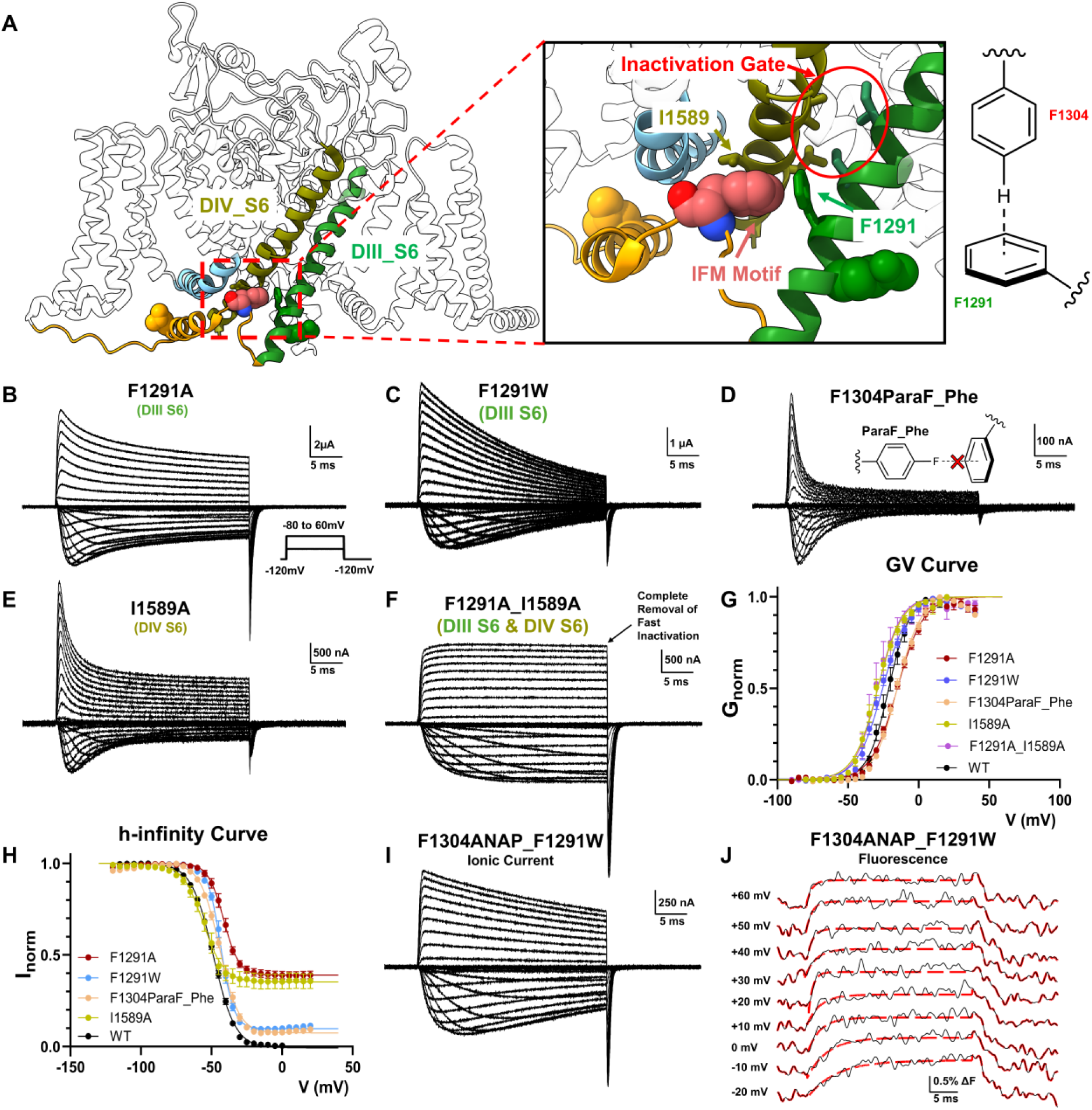
F1291 and I1589 relay IFM movement to the pore through hydrophobic and aromatic interactions. **A)** Interactions of IFM motif (pink) with F1291 in DIII S6 (green) and I1589 in DIV S6 (olive). Previously identified fast inactivate is highlighted in red circle. Relative positions between IFM and F1291 suggest a potential T-shaped π-π interaction. **B) to E)** Example of ionic current from F1291 and I1589 mutants. In F1291A, F1291W and I1589A, significant fast inactivation removal was observed. Intriguingly, replacement of a single hydrogen atom to fluorine at the para position of F1304 also led to incomplete inactivation, suggesting the existence of the suspected aromatic interaction. Inset shows voltage protocol **F)** Double mutant of F1291A_I1589A led to complete removal of fast inactivation. No significant current decay was observed after 30ms of depolarization. **G)** GV curves of the all the mutants shown previously. N ≥ 4. **H)** h-infinity curves for the mutants shown previously. N ≥ 4. **I)** Example of Ionic current from F1304ANAP_F1291W. Fast inactivation was largely absent. **J)** Fluorescence signal from F1304ANAP_F1291W fitted with one exponential component. Different from F1304ANAP, fluorescence signals from F1304_F1291W can be adequately described by a mono exponential. By disrupting the interactions of IFM motif with the F1291, the second component of the fluorescence signal seen in F1304ANAP is eliminated. Data is shown as mean ± SEM.

In in DIII S6, when F1291 is mutated to either smaller or bigger residues, significant amount of fast inactivation was removed (Fig. 4B, C and H), suggesting that the precise size of the side chain is important for this interaction. Additionally, structural analysis ^33^ also pointed out the possibility of a T-shaped π-π interaction between the phenylalanine in the IFM motif and F1291 (Fig. 4A). To precisely target this potential aromatic interaction, we utilized another unnatural amino acid, 4-fluorophenylalanine (ParaF-Phe). Substitution of the hydrogen at the para position on the aromatic ring targets the positive charge on the edge of the aromatic ring and would be predicted to weaken a T-shaped π-π interaction with a minimal impact on the volume of side chain ^34^. In support of this notion, substitution of only one atom in F1304 via encoding of ParaF-Phe led to not only slowed, but also incomplete fast inactivation and faster recovery from inactivation (Supple. Fig. 3), suggesting a functional role for this π-π interaction F1304 and F1291 (Fig. 4D, H).

Mutagenesis in DIV S6 revealed that changing I1589 led to similar phenotype observed by mutations in DIII S6. I1589A severely impeded fast inactivation and showed large non-inactivating steady-state current (Fig. 4E, H). As the inactivation gate is made of large hydrophobic residues in S6 of both domain III and IV, the combination of F1291A in S6-DIII and I1589A in S6-DIV should eliminate fast inactivation. Indeed, combining both mutations in a single construct, F1291A_I1589A, produced channels where fast inactivation was completely removed. No significant reduction in current amplitude was seen after 30ms of depolarization (Fig. 4F).

Our mutagenesis experiments on F1291 and I1589 confirmed their importance for fast inactivation. However, these results did not directly demonstrate those are residues that coupled the IFM motif to the fast inactivation gate at the pore. To provide direct evidence, we investigated the movement of the IFM motif in the absence of the identified interactions. We recorded the ionic current and fluorescence signal in F1304ANAP_F1291W (Fig. 4I, J). Fast inactivation was largely removed in F1304ANAP_F1291W, but still, clear fluorescence signals were resolved from the IFM position. Evidently, disrupting the interaction between F1291 and F1304 did not abolish the binding of the IFM motif. The kinetics of the fluorescence signals were significantly faster than the residual fast inactivation. These observations are consistent with our previous results suggesting that the fast component of the IFM movement was triggered by the DIV VSD and coupled by the hydrogen bonds, instead of interactions in the hydrophobic pocket. More importantly, now with disrupted interaction with F1291, the IFM motif movement showed only one fast fluorescence component and the second slow movement seen before in F1304ANAP alone was largely absent (Fig. 4J), providing direct evidence that the identified residues are responsible for coupling the IFM motif and the pore.

### Fast Inactivation Process Requires Sequential Conformational Changes

Finally, we monitored the movement of the pore during fast inactivation by encoding ANAP at Q1293, a position at the bottom of the pore-lining DIII S6 segment (Fig. 5A). Incorporation of ANAP at Q1293 led to minimal alteration in fast inactivation (Fig. 5B, D) and activation (Fig. 5C). The fluorescence signal from Q1293ANAP mirrored the development of fast inactivation, suggesting the movement seen at this part of the S6 helix reflects mostly conformational changes associated with fast inactivation instead of activation (Fig. 5E). Interestingly, there was a ∼1ms delay to the onset of the fast-inactivation-associated fluorescence signal. This delay is similar to the previous observation in pronase treated giant squid axon^35^ (Fig. 5B inset). The observed delay also resembles the classical Cole-Moore shift ^36^ and indicates that additional conformational changes are required before channel can enter into the fast inactivated state. It is likely that the two movements seen at the IFM position happen in a sequential manner in that the DIII_DIV linker needs to move to posit the IFM at the right location before the second, slower conformation change can happen and triggers the fast inactivation. Supporting this notion, in the case of L1319ANAP_F1291A (Fig. 5F), the fluorescence signal was unaltered by abolishing the second, slower interaction (Fig. 5G), suggesting the DIII_DIV linker movement happened upstream of the second interaction and the fast inactivation process is best described as sequential conformational changes.

**Figure 5:**
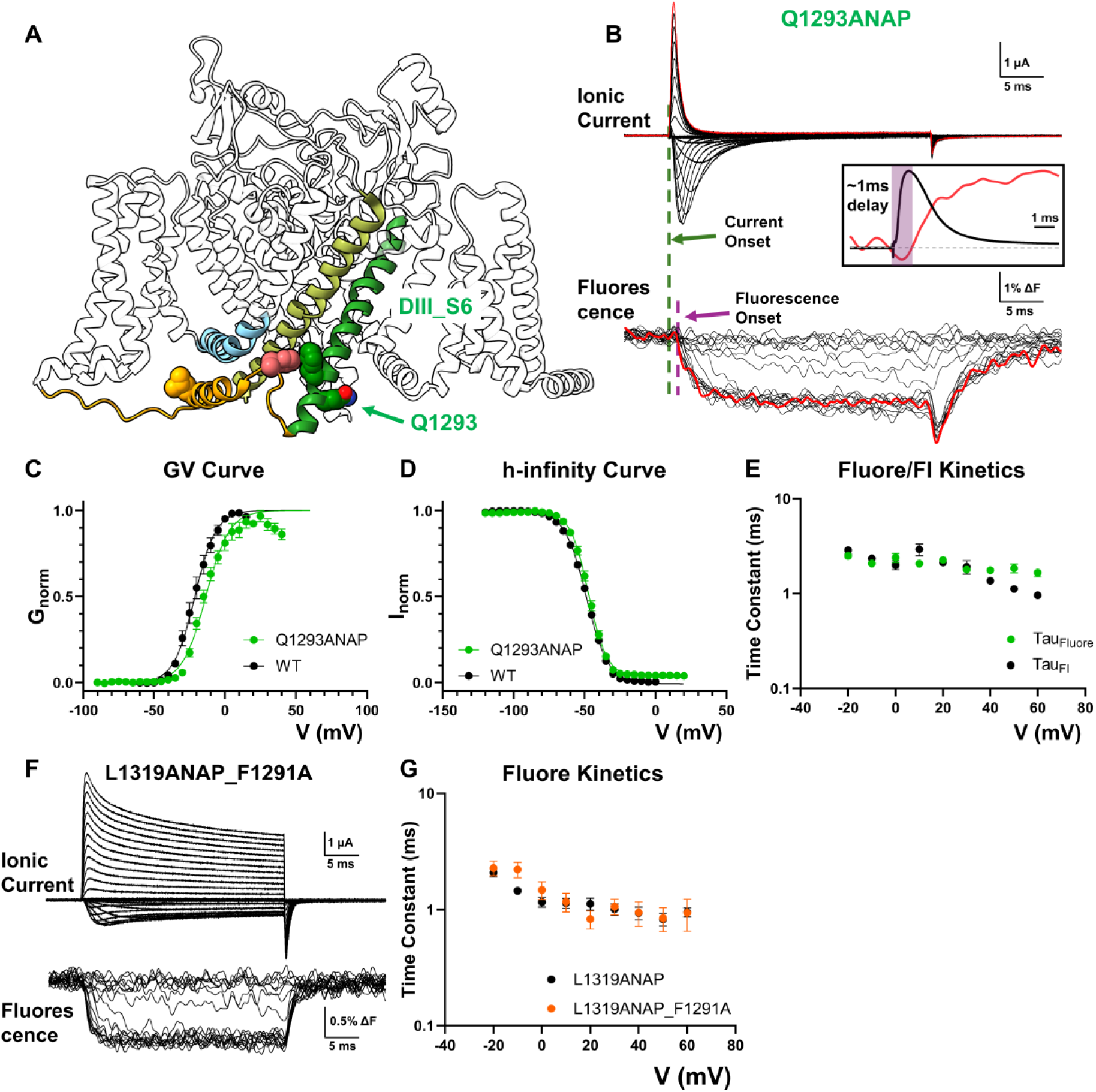
Pore movement as the last conformational step in fast inactivation. **A)** To track the pore movement during fast inactivation, ANAP was incorporated at Q1293 position at the bottom of DIII S6 (green). **B)** Example of ionic current and fluorescence signal recorded from Q1293ANAP. Inset shows overlapped comparison of ionic current and fluorescence signal at 60mV. There is a ∼1ms delay to the onset of fluorescence signal, suggesting a sequential movement for fast inactivation. **C), D)** GV curve and h-infinity curve for Q1293ANAP. No drastic difference was observed. N = 5 for Q1293ANAP. **E)** Comparison of fluorescence kinetics and fast inactivation kinetics. Fluorescence kinetics follows closely the fast inactivation development, suggesting the conformational change observed at this position reports mostly pore movement associated with fast inactivation. N = 10. **F)** Example of ionic current and fluorescence signal from L1319ANAP_F1291A. **G)** Comparison of fluorescence kinetics between L1319ANAP and L1319ANAP_F1291A. No significant difference in kinetics is seen in the DIII_DIV linker movement after abolishing the interaction between IFM and F1291, favoring a sequential model for fast inactivation. N = 7 for L1319ANAP_F1291A and N = 8 for L1319ANAP. Data is shown as mean ± SEM.

## Discussion

### The IFM Motif Couples the DIV_VSD and Fast Inactivation Gate

Connections between VSD movements and fast inactivation has long been established. From the pronase perfused giant squid axon ^35^ to charge neutralization experiments ^32^ on DIV VSD, it is clear that VSD movements are required for the fast inactivation. On the other hand, fast inactivation could also influence the VSD movements, as was shown in charge immobilization experiments ^10,37^. Our results indicate that the IFM motif serves as a structural coupler that connects the voltage-driven movement of the DIV VSD to the inner mouth of the pore that inactivates sodium channels. By positioning ANAP into the IFM motif, we find that the resulting fluorescence signal reveals two sequential movements at IFM motif during fast inactivation. The fast movement originates from the DIII_DIV linker movement, coupled to the DIV VSD movement and the slower movement directly triggers the closure of the fast inactivation. Evidently, the movement of IFM motif transduces the activation of DIV VSD directly to closure of the fast inactivation gate at the pore. These two movements also explain the domain specific charge immobilization during fast inactivation shown previously in MTS modification experiments ^10,38^. It is likely that the movement of the DIII_DIV linker triggers the immobilization of DIV VSD by binding of the IFM with the DIV S4-S5 linker, while the closure of the fast inactivation gate immobilizes DIII VSD.

### Functional Identification and Validation of the Binding Site for IFM motif

In this work, we further elucidated a network of hydrogen bonds that forms the structural connection between DIV VSD and cytoplasmic DIII_DIV linker. One intriguing observation is that in both hydrogen bonds, it is the carbonyl groups in the peptide backbone which serve as the H-bond acceptors. In particular, N1477 at DIV S4_S5 linker forms a direct interaction with the backbone carbonyl group of the phenylalanine in IFM motif. One natural conclusion from this observation would be that the exact identify of the residue at the second position of IFM motif shouldn’t influence the formation of the hydrogen bonds and subsequently shouldn’t impede the “binding” of IFM motif. The binding of IFM motif has been traditionally associated with interactions between the side chain of the phenylalanine residue and its surrounding hydrophobic pocket. However, our results argue that the driving force of the IFM motif binding comes mostly from the hydrogen bond formation with the peptide backbones. In other words, the binding site of the IFM motif is formed not by the hydrophobic pocket but rather Q1309 in DIII_DIV linker and N1477 in DIV S4_S5 linker. It has been observed in molecular dynamic simulations before that mutation N1662D, a disease causing mutant in hNav1.2 (equivalent as N1477 in rNav1.4), led to significantly decreased dwell time of IFM in the hydrophobic pocket ^39^. This is consistent with our electrophysiological results. Removing one of the hydrogen bonds in Q1309A led to an unstable fast inactivated state as was shown by the increased recovery speed from inactivation. The fluorescence results, which arguably track motions in this area, also support this conclusion. Disruption of the hydrophobic pocket at F1291, while resulting in significant removal of fast inactivation (F1291W), didn’t impede the binding of IFM motif in F1304ANAP_F1291W (Fig. 5F). On the other hand, a mild mutation Q1309A clearly led to impaired coupling between DIV VSD and DIII_DIV linker as was seen in L1319ANAP_Q1309A. Finally, charge immobilization experiments in IQM also demonstrated that despite almost complete removal of fast inactivation from ionic current, ∼30% (in contrast to 60% in WT) of total gating charge still became immobilized after 24ms of depolarization (Supple. Fig. 4). Clearly, even in IQM channels, some conformational changes along the fast inactivation pathway are still able to occur. It is likely the DIII_DIV linker in this case still binds to the identified residues, leading to some degree of charge immobilization, likely only DIV VSD^13,38^, but as no further movement triggers the closure of the fast inactivation gate, the immobilization of DIII does not occur. Taken together, the present data strongly favors N1477 and P1473 in DIV S4_S5 linker as the binding sites for IFM motif, instead of a hydrophobic pocket.

### Molecular Mechanism of Fast Inactivation

The current and available data are therefore consistent with a “lock and key” model for fast inactivation in voltage-gated sodium channels. Figure 6 is a schematic representation of the sequence of events that lead to fast inactivation. Upon depolarization, the sensors of domain I, II and III activate and the channel opens (fig. 6A, E, I). Normally, domain IV sensor activates later ^30,32^ and upon its activation it exposes the binding site for IFM motif at P1470 and N1477 in DIV S4_S5 linker, similar to a keyhole becoming available (Fig. 6 B, F, J). The IFM-containing DIII_DIV linker, likely mobile in aqueous solution, binds to DIV S4_S5 linker via the H-bonds (Fig. 6 C, G) and posits IFM motif, the key, into the hydrophobic pocket, the keyhole (Fig. 6K). The binding of IFM motif allows the interactions with F1291 and I1589 through hydrophobic and aromatic interactions to occur, leading to a second conformational change at the hydrophobic pocket. In F1291Q, despite some steady state current, the majority of fast inactivation persists, dissimilar to F1304Q (IQM) phenotype (Supple. Fig. 5). This asymmetric effect suggests that for interactions between F1291 and I1589 to occur, another conformational change must occur. Most likely, DIII S6 and DIV S6 helices undergo a rotational movement to position F291 and I1589 into the hydrophobic pocket interacting with the IFM motif and thus locking the two S6 helices in position. At the same time, the rotations position the four large hydrophobic amino acids forming the inactivation gate into the conduction pathway interrupting the flow of Na+ ions, thus producing fast inactivation (Fig. 6D, H, L). Modifications to either IFM motif or the hydrophobic pocket would lead to incompatibility between the key and the lock, leading to impaired inactivation.

**Figure 6:**
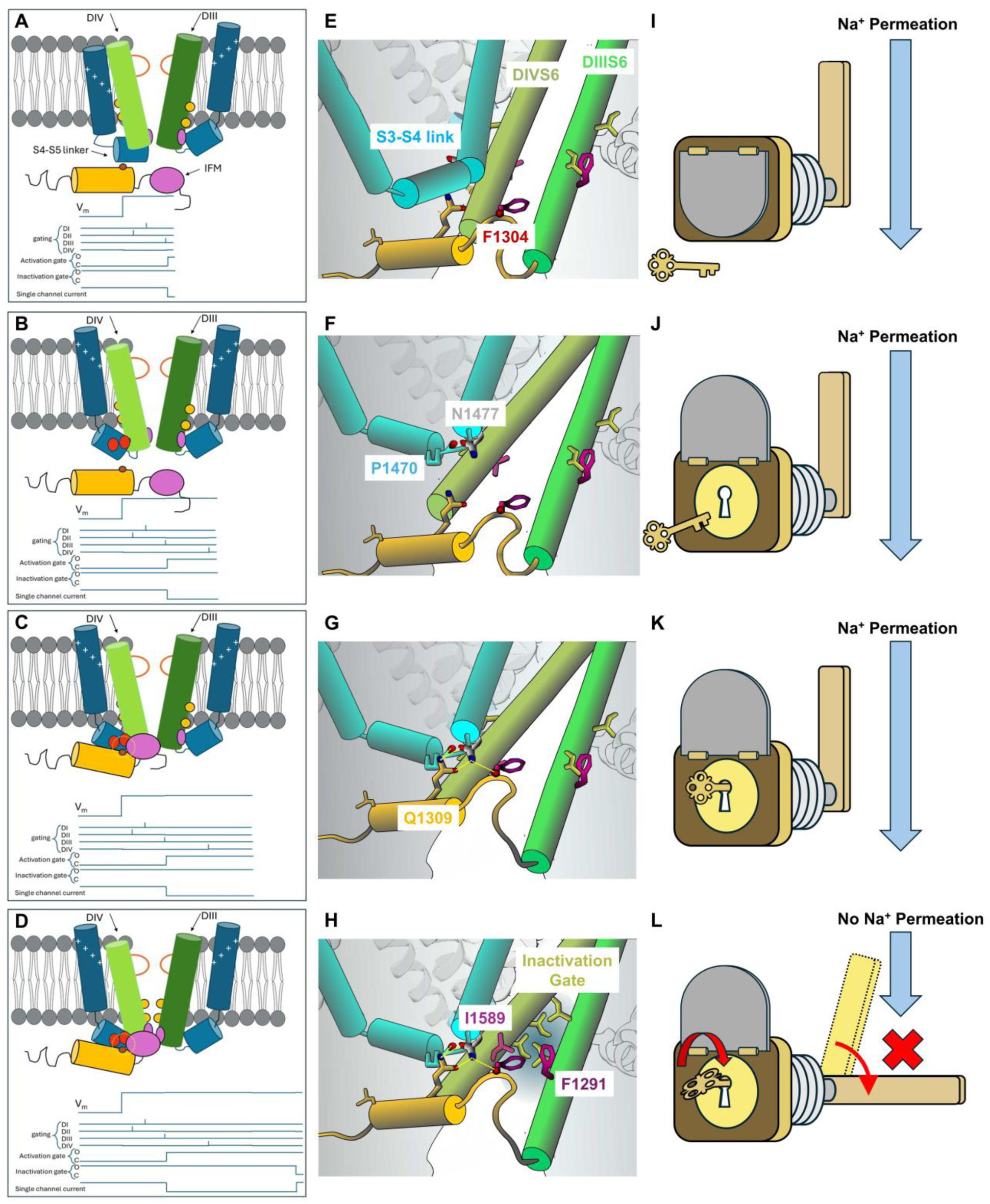
Conformational changes and the “lock and key” model for fast inactivation. **Left** column is a schematic diagram of the sequence of events. The **Middle** column shows details of the residues involved. The **Right** column shows the lock and key analogy. Upon depolarization, VSD from DI to DIII activates and subsequently opens the channel. Given that DIV VSD resides in the resting state (**A** and **E**), the hydrogen bond mediated binding pocket for IFM motif is shielded (the lock is covered, **I**). Maintaining the depolarization, it leads to the activation DIV VSD which reorganizes the DIV S4_S5 linker and displays P1470 and N1477 towards the intracellular aqueous environment (**B** and **F**). This process is similar to opening a guard on the lock and exposing the keyhole (**J**). Now the DIII_DIV linker, possibly mobile in the aqueous environment, binds to the DIV S4_S5 linker positioning the IFM motif, the key, into the hydrophobic pocket (**C** and **G**), into the keyhole (**K**). The binding of IFM motif allows the interactions of F1291 and I1589 with the phenylalanine in IFM motif, creating a rotation of the pore-forming S6 helices in DIII and DIV (**D** and **H**), like turning the key in the keyhole (**L**). This helical rotation in turn locates the hydrophobic gate (yellow residues in **D** and **H** and blocking gate in **L**) in the permeation pathway, blocking the Na^+^ permeation as a result.

Recovery from inactivation requires disengagement of the IFM motif. Results of F1304ANAP demonstrate that the recovery from inactivation and off-fluorescence signal share similar time course, supporting this idea (Supple. Fig. 1). The two major conformational changes that occurred during fast inactivation are responsible for immobilization of DIV VSD and DIII VSD respectively as suggested by the IQM gating current result ^13,37^ (Supple. Fig. 4). Under hyperpolarized voltages, a transient unbinding of the IFM motif would allow the deactivation of voltage sensors in DIII and DIV that are previously immobilized. This inward movement of the sensors subsequently abolish the binding site for IFM motif, similar to covering the keyhole on the lock and allows the channels to recover from the fast inactivated state.

### Alternative Fast Inactivated State in Nav Channels

Here we addressed the molecular pathway from open channels to the fast inactivated state. It is important to point out that closed Nav channels can also inactivate, bypassing the open state altogether, and this is essential for physiological processes ^10,40,41^. This closed-state inactivation, or steady-state inactivation, prevents Na^+^ conduction upon hyperpolarization and the recovery from closed-state inactivation determines the absolute refractory period in firing neurons. Structural work suggests a similar pore conformation in the closed inactivated state and open inactivated state ^19,33^. Fluorescent measurements on individual sensor movements have demonstrated that DIV VSD can activate at more hyperpolarized voltages compared to the other VSDs ^30^. Then, as the movement of DIV VSD alone seems to be enough to trigger inactivation while opening requires all the other VSDs to be activated, it is possible that fast inactivation may occur regardless of an open or a closed pore. When we monitored the pore movement at Q1293 position, two components of fluorescence signals were observed upon hyperpolarization (Fig. 5B). It is possible these two components capture the transitions from open inactivated state to closed inactivated state and from closed inactivated state to closed state respectively. However, further investigations are required to elucidate the exact structural rearrangement at the pore during these transitions. But most likely, the observed sequential conformational changes also occur during closed state inactivation.

### Evolutionary and Isoform Conservation of the Identified Fast Inactivation Apparatus

We compared Nav channels from Bilateria, Cnidaria and Choanoflagelleta (Supple. Fig. 5) and intriguingly found the newly identified residues are highly conserved, some even more conserved than the IFM motif itself. F1291, N1477 and Q1309 are conserved across all sequences compared, even in the ancestral form of Nav channel in *Monosiga brevicollis* ^42^. In the case of purple sea urchin, *Strongylocentrotus purpuratus*, an isoleucine replaces the phenylalanine in the IFM motif but nevertheless, all three residues remain conserved at the equivalent positions. I1589, on the other hand, is less conserved. In the ancestral Nav channel, a valine is found at 1589 equivalent position and valine is the only alternative amino acid found at this position across different phyla. The high conservation of the newly identified fast inactivation components strongly supports the idea that the molecular mechanism described here in mammalian Nav1.4 channel is conserved across the animal kingdom and appears at the same time as the evolutionary invention of the Nav channel in Choanoflagelleta.

## Material and Method

### Site-directed mutagenesis and cRNA *in-vitro* synthesis

rNav1.4 in pBSTA vector, flanked by β-globin sequences, was used in this study for all the physiological experiments. Point mutations were generated using QuikChange^TM^ mutagenesis method (Agilent). The PCR products were first digested by DpnI and were used to transform the XL10-gold ultra-competent cells (Agilent). After ampicillin resistance screening, plasmids were purified from the colonies using standard miniprep protocols. Purified plasmids were sent to Plasmidsaurus for whole plasmid sequencing to confirm the introduction of the mutations. Verified plasmids were first linearized and later *in vitro* transcribed to cRNA with mMESSAGE mMACHINE™ T7 Transcription Kit (Thermo Fisher Scientific).

### *Xenopus laevis* oocyte preparation and channel expression

Ovaries of *Xenopus laevis* were purchased from XENOPUS1 (Dexter, Michigan). The follicular membrane was removed using collagenase type II (Worthington Biochemical Corporation) 2 mg/mL with bovine serum albumin at 1mg/mL (BSA). After defolliculation, stage V–VI oocytes were then selected and microinjected with 50-100 ng cRNA. Injected oocytes were incubated at 18 °C for 1-5 days in SOS solution (in mM: 100 NaCl, 5 KCl, 2 CaCl^2^, 0.1 EDTA, and 10 HEPES at pH 7.4) supplemented with 50 µg/mL. Unless otherwise stated, all chemicals were purchased from Sigma-Aldrich.

### Cut-open voltage clamp on *Xenopus laevis* oocytes

Macroscopic ionic and gating current were recorded using cut-open voltage clamp technique^43^. Micropipettes filled with 3M CsCl, with resistance between 0.6 and 1.2 MΩ were used to measure the internal voltage of the oocytes. Current data were first filtered online at 20 kHz with a low-pass 4 pole Bessel filter and sampled by a 16-bit A/D (USB-1604; Measurement Computing, Norton, MA) converter at 1 MHz. A feedbackback Peltier device was used to control the temperature at 13 ± 1 °C for all experiments. For ionic current experiments, unless otherwise stated, were conducted in external solution consisted of in mM: 28 Na methylsulfonate (MES), 92 N-methyl-D-glucamine (NMG) MES, 2 Ca MES, 10 HEPES, 0.1 EDTA, pH = 7.4 and internal solution consisted of in mM: 12 Na MES, 108 NMG MES, 10 HEPES, 2 EGTA, pH = 7.4. The capacitive transient was manually compensated with a dedicated circuit and further removed by an online P/-4 protocol with a holding voltage of either -80 or -90 mV^10^. For gating current experiments, all experiments were conducted in external solutions consisted of in mM: 120 NMG MES, 2 Ca MES, 10 HEPES, 0.1 EDTA, 750nM TTX pH = 7.4 and in internal solution consisted of in mM: 120 NMG MES, 10 HEPES, 2 EGTA, pH = 7.4. In this case, an online P/4 at 40mV was used to subtract the capacitive transients. Voltage clamp speed measured by capacitive transient current gave a time constant around 70 µs.

### Unnatural Amino Acid Incorporation

Two methods were used to incorporate the unnatural amino acid in Nav channels. For ANAP, two separate cytoplasm injection were performed. First injection introduced into oocytes four different components: ANAP amber suppression tRNA at 2 ng/nL, ANAP synthetase at 0.3 ng/nL, mutated *Xenopus laevis* release factor 1 at 0.3 ng/nL and the unnatural amino acid ANAP itself in methyl ester form (Cayman Chemical) at 2mM. 24 hours after the first injection, channel cRNA bearing the amber stop codon mutation at the desired location was injected at 1 ng/nL. 24 to 72 hours after the second injection, the oocytes were used in voltage clamp fluorimetry experiments.

4-fluoro-L-phenylalanine was appended to pyl tRNA as previously reported^44,45^.Briefly, the cyanomethyl ester of N-Boc-4-fluoro-L-Phe was reacted with the dinucleotide pdCpA in dimethylformamide, purified on HPLC and deprotected with trifluoroacetic acid. A 3 mM stock solution on the amino acid-dinucleotide chimera in dimethylsulfoxide was prepared and added to T4 RNA ligase and pyl tRNA lacking the terminal CA nucleotides to assemble the full-length tRNA with 4-fluoro-L-Phe attached. Pellets (1 nmol) were stored dry at -78°C until needed. Lyophilized conjugated tRNA was resuspended in 3mM Na Acetate at pH 5.5 to an approximate concentration of 2.5 µg/µL. Conjugated tRNA and cRNA of the mutated channel were introduced simultaneously into the oocytes and after 24 to 48 hours of expression, oocytes were used in voltage-clamp experiments.

### Voltage Clamp Fluorimetry Experiments

An avalanche photodiode or a photomultiplier was used to record the fluorescence signal. A 365nM LED (Thor Lab) was used as the excitation light source. The filter set used in the study was described previously^46^. In the case of the photomultiplier (H10304-20-NN, Hamamatsu, Japan), the output signal was first processed by a home-built I to V converter, filtered by a low-pass 8 pole Bessel filter at 10kHz and sampled by the AD converter every 10 µs. Photobleaching was later subtracted with a Matlab routine and then digitally filtered at 5kHz or 1kHz.

### Data Analysis

Data recorded in the work is analyzed by GraphPad 9 (Prism), Excel (Microsoft), Matlab R2022a (Mathworks), and in-house software (Analysis and GPatchM).

**i)** GV curves calculation: Ionic conductance was calculated by dividing the peak ionic current by the driving force determined experimentally. After normalization, GV curves were fitted with a two-state model described below,

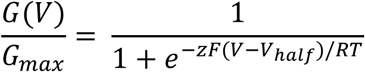

, where R is the ideal gas constant, T is the absolute temperature in Kelvin, z is the apparent charge, F is the Faraday constant.

**ii)** FV curve calculation: Steady state fluorescence signal was normalized to maximum fluorescence and then described with the model below,

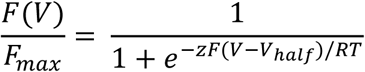

, where R, T, F and z are as previously defined

**iii)** Steady state fast inactivation curve (*h*-infinity curve) was calculated by plotting normalized peak Na^+^ current during test pulse after 50ms conditioning pulse voltage and fitted using a two-state model,

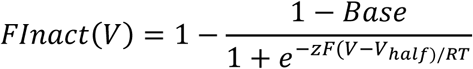

, where Base is the steady state current, and R, T, F and z are as previously defined.

**iv)** The time constant of fast inactivation, gating charge movement or fluorescence kinetics were calculated by fitting the signals with either one or two exponential decays using the general equation below,

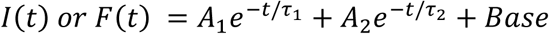

, where *A*_1_, *τ*_1_, *A*_2_, and *τ*_2_ represent amplitudes and time constants of the first and second components, respectively. Base represents the steady state current or fluorescence. When one exponential was used, only *A*_1_, and *τ*_1_ were fitted to the data.

**v)** Recovery from fast inactivation was calculated by dividing the peak current in the control pulses by the peak current during test pulses. The data was then described by either one or two component association model,

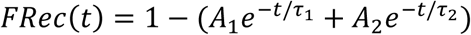

, where *A*_1_, *τ*_1_, *A*_2_, and *τ*_2_ represent amplitudes and time constants of the first and second components, respectively. When one exponential was used, only *A*_1_, and *τ*_1_ were fitted to the data.

## Acknowledgements

We would like to thank Gethiely Gasparini and Hlafira Polishchuk for oocyte preparation, DNA mutation and RNA preparation. YL and FB are supported by NIH grant R01-GM030376, R01-GM150272 and National Science Foundation Award QuBBE QLCI (NSF OMA-2121044). CAA and JDG are supported by R35GM148239.

## Data Availability

All data and resources presented in the paper will be made available upon reasonable request. The raw data used to generate the figures are submitted as Source Data.

## Author Contribution

YL performed electrophysiological and voltage fluorimetry experiments.

JDG performed the tRNA ligation experiments

YL analyzed the data.

YL and FB conceptualized the project.

YL, CAA and FB interpreted the data.

YL wrote the original draft.

YL, JDG, CAA and FB edited and reviewed the paper

C.A. and F.B. supervised the project and oversaw funding acquisition.

## Supplementary Tables

**Supplementary Table 1:** GV curves fittings of all the mutants studied in this work. For simplicity’s sake, only best fit values are shown.

|  | **V50 (mV)** | **z** | **N** |
| --- | --- | --- | --- |
| **WT** | -21.19 | 3.32 | 4 |
| **Q1309A** | -18.59 | 2.96 | 5 |
| **Q1309L** | -20.90 | 2.61 | 4 |
| **N1477D** | -22.64 | 2.52 | 5 |
| **I1589A** | -29.01 | 2.86 | 5 |
| **F1291A** | 15.09 | 2.65 | 5 |
| **F1291W** | -25.52 | 2.74 | 7 |
| **F1291A_I1589A** | -28.95 | 3.04 | 4 |
| **F1291ParaF_Phe** | -15.14 | 2.96 | 5 |
| **Q1293ANAP** | -13.41 | 2.97 | 5 |
| **F1304ANAP** | -21.37 | 3.63 | 5 |
| **L1319ANAP** | -18.12 | 2.77 | 5 |
| **L1319ANAP_F1291A** | -28.81 | 2.77 | 5 |
| **L1319ANAP_Q1309A** | -21.29 | 3.33 | 5 |

**Supplementary Table 2:** hInfinity curves fittings of all the inactivating mutants studied in this work. For simplicity’s sake, only best fit values are shown.

|  | **V50 (mV)** | **z** | **Base** | **N** |
| --- | --- | --- | --- | --- |
| **WT** | -48.74 | 3.19 | -0.74% | 4 |
| **F1304ANAP** | -40.28 | 5.2 | 3.35% | 5 |
| **L1319ANAP** | -57.13 | 3.07 | 0.36% | 6 |
| **Q1309A** | -41.74 | 4.55 | 6.61% | 4 |
| **Q1309L** | -45.34 | 4.92 | 26.53% | 4 |
| **N1477D** | -33.57 | 6.17 | 29.04% | 6 |
| **I1589A** | -55.95 | 3.34 | 35.19% | 5 |
| **F1304ParaF_Phe** | -44.34 | 3.62 | 7.38% | 5 |
| **F1291A** | -41.44 | 4.51 | 38.89% | 5 |
| **F1291W** | -43.5 | 5.27 | 9.64% | 7 |
| **Q1293ANAP** | -47.33 | 3.81 | 3.59% | 5 |

**Supplementary Table 2:** Best fitting parameters for the FV curves shown in the figures.

|  | **V50 (mV)** | **z** | **N** |
| --- | --- | --- | --- |
| **F1304ANAP** | -36.44 | 3.25 | 5 |
| **L1319ANAP** | -63.85 | 1.4 | 10 |

**Supplementary Figure 1:**
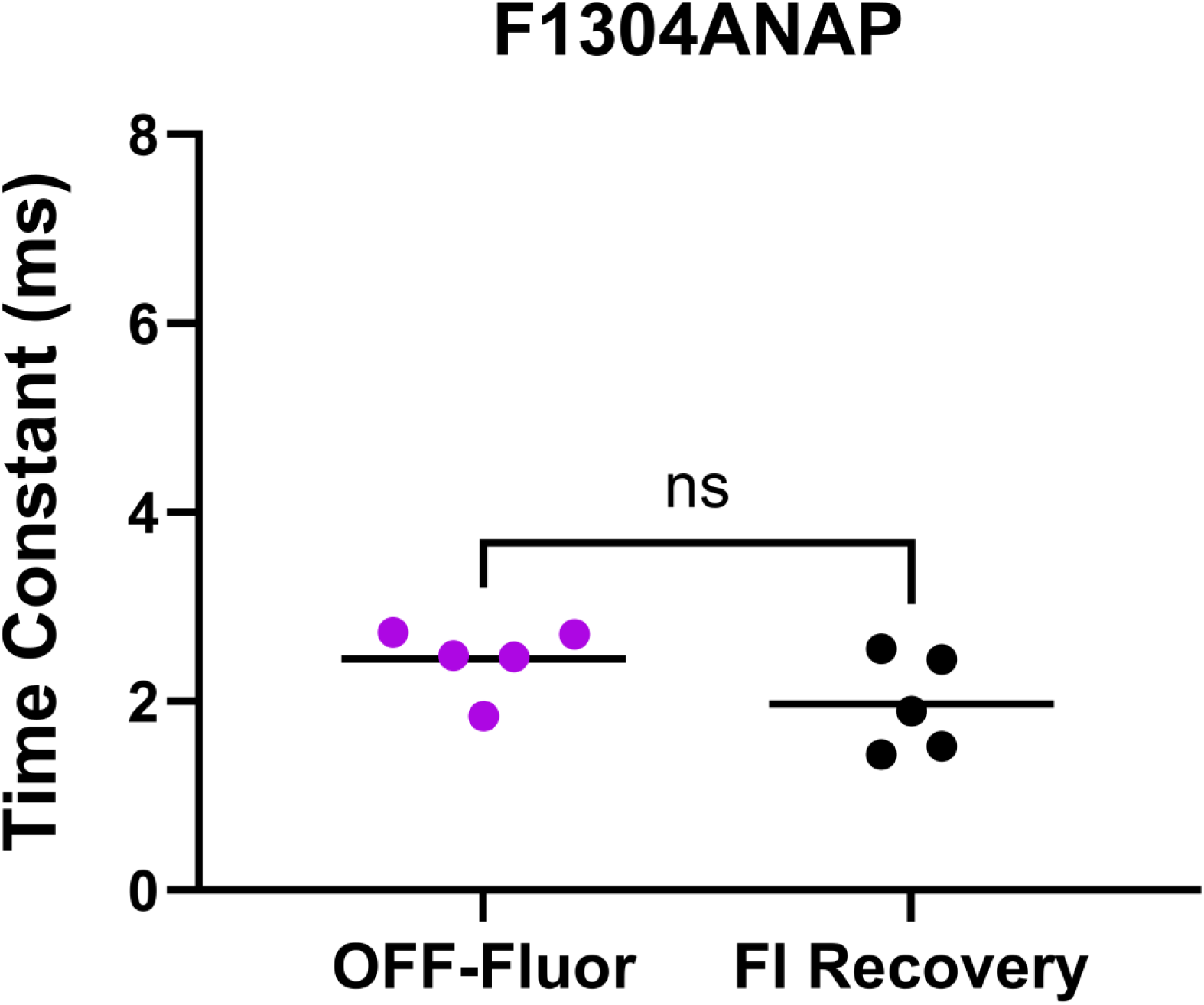
Comparison of off time constants of fluorescence signal (OFF-Fluor) and recovery from fast inactivation (FI Recovery) in F1304ANAP at - 120mV. Their kinetics are not significantly different suggesting the unbinding of IFM motif tracks the recovery from fast inactivation

**Supplementary Figure 2:**
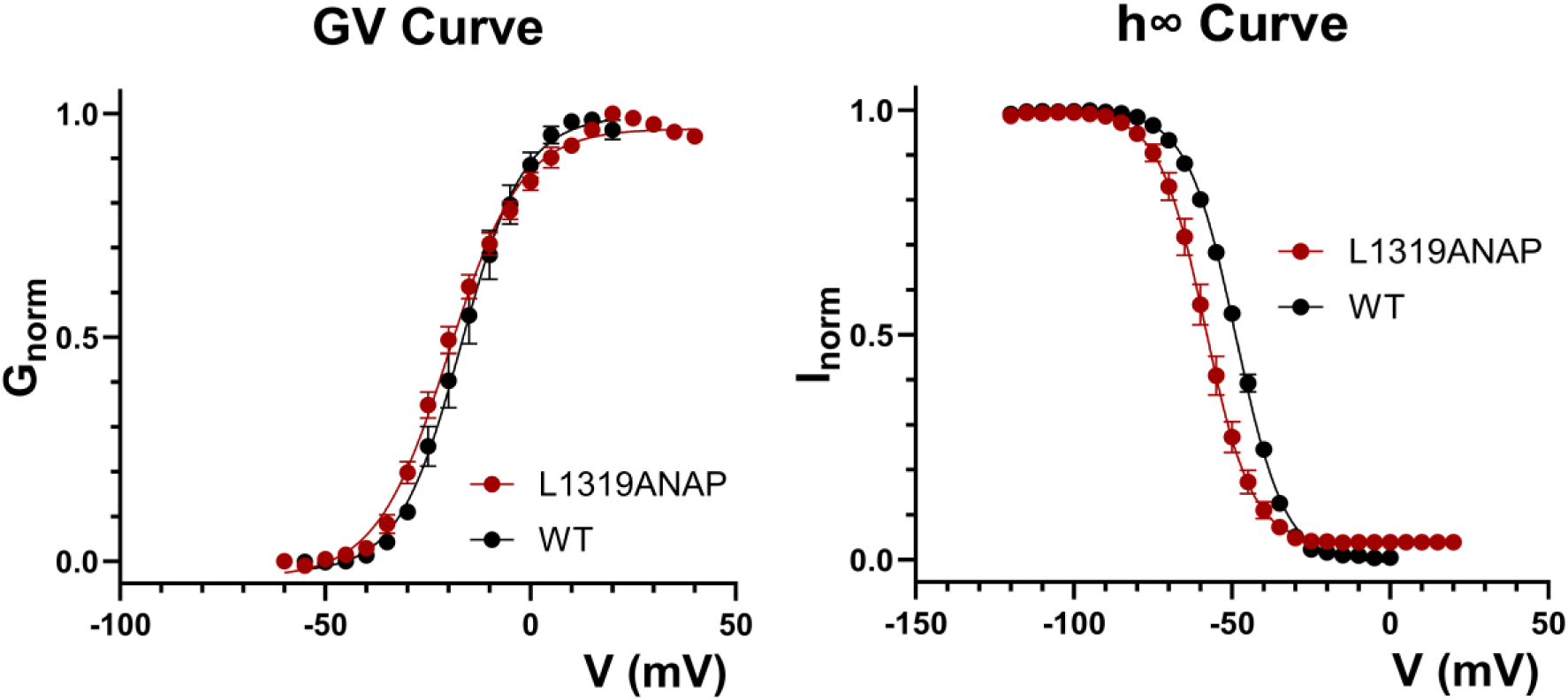
GV curve and h-infinity curve of L1319ANAP. Incorporation of ANAP at L1319 position doesn’t disrupt activation and only mildly shifted the h-infinity curve to the left. N = 4 for WT, N = 5 for L1319ANAP.

**Supplementary Figure 3:**
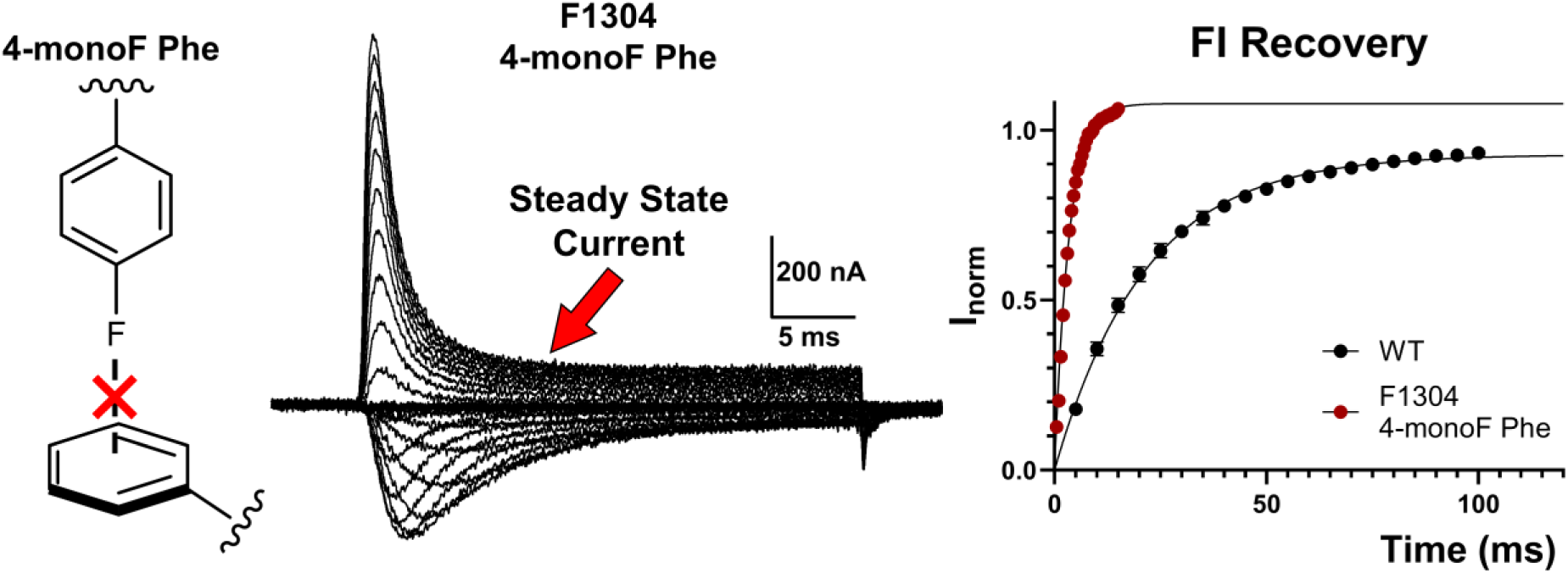
Fast inactivation recovery kinetics of F1304 4-monoF Phe. The recovery from fast inactivation is significantly faster than the WT, suggesting a destabilized inactivated state.

**Supplementary Figure 4:**
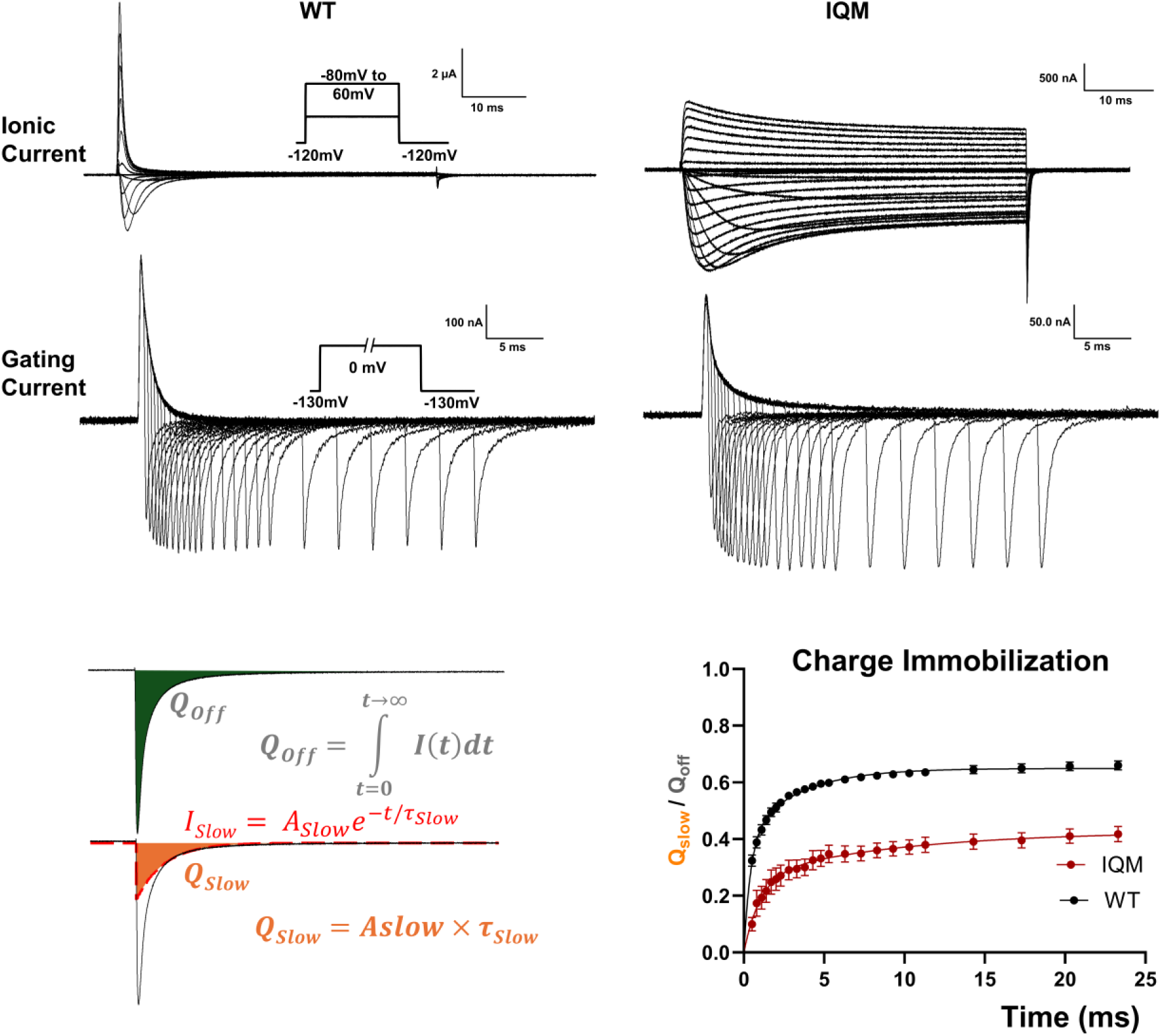
Comparison of gating charge immobilization in WT and IQM. Despite fast inactivation is largely removed in IQM, 30% of gating charge is still immobilized after 24ms of depolarization.

**Supplementary Figure 5:**
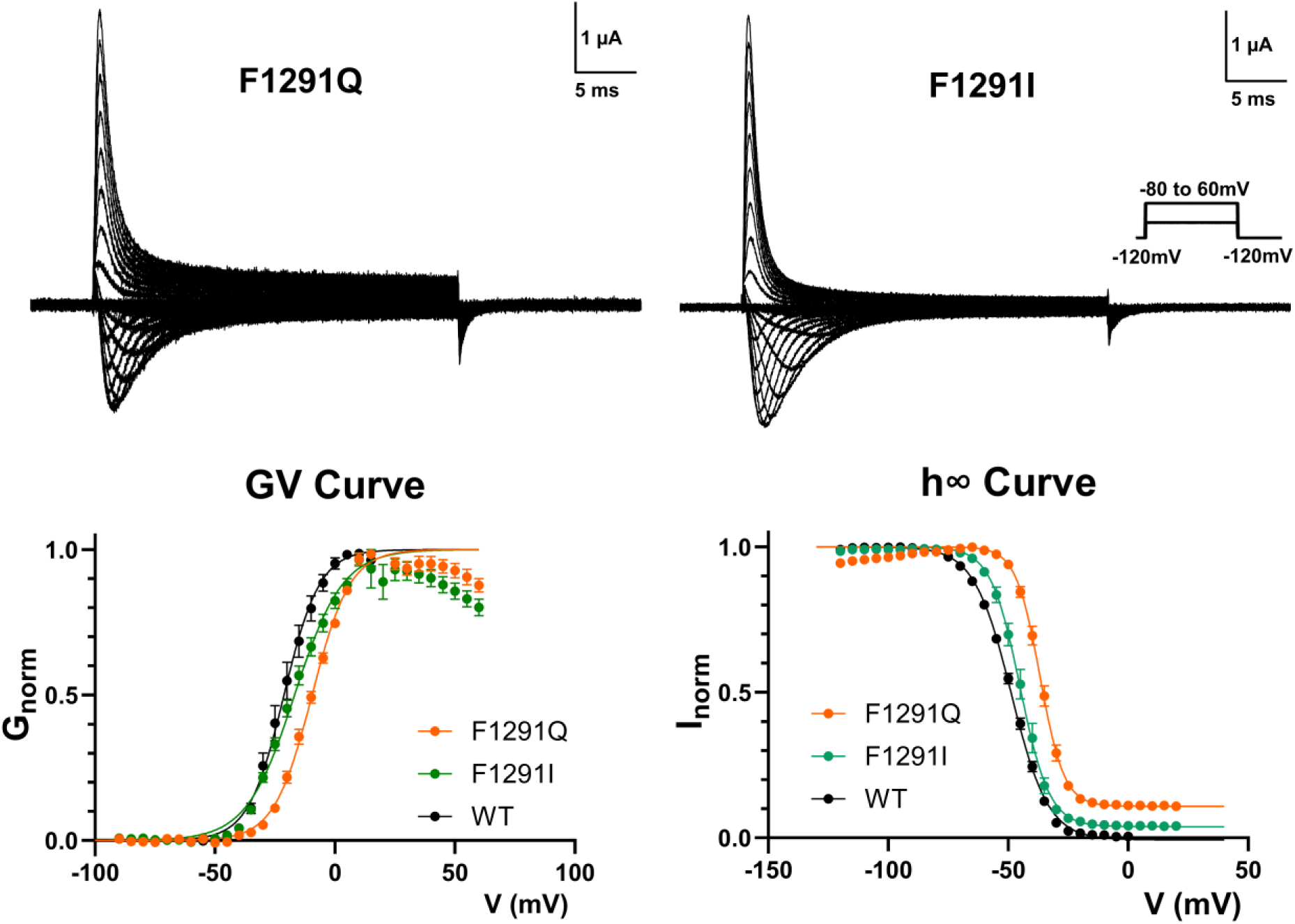
Other mutants tested at F1291 position, F1291Q and F1291I and their GV, h-infinity curve.

**Supplementary Figure 6:**
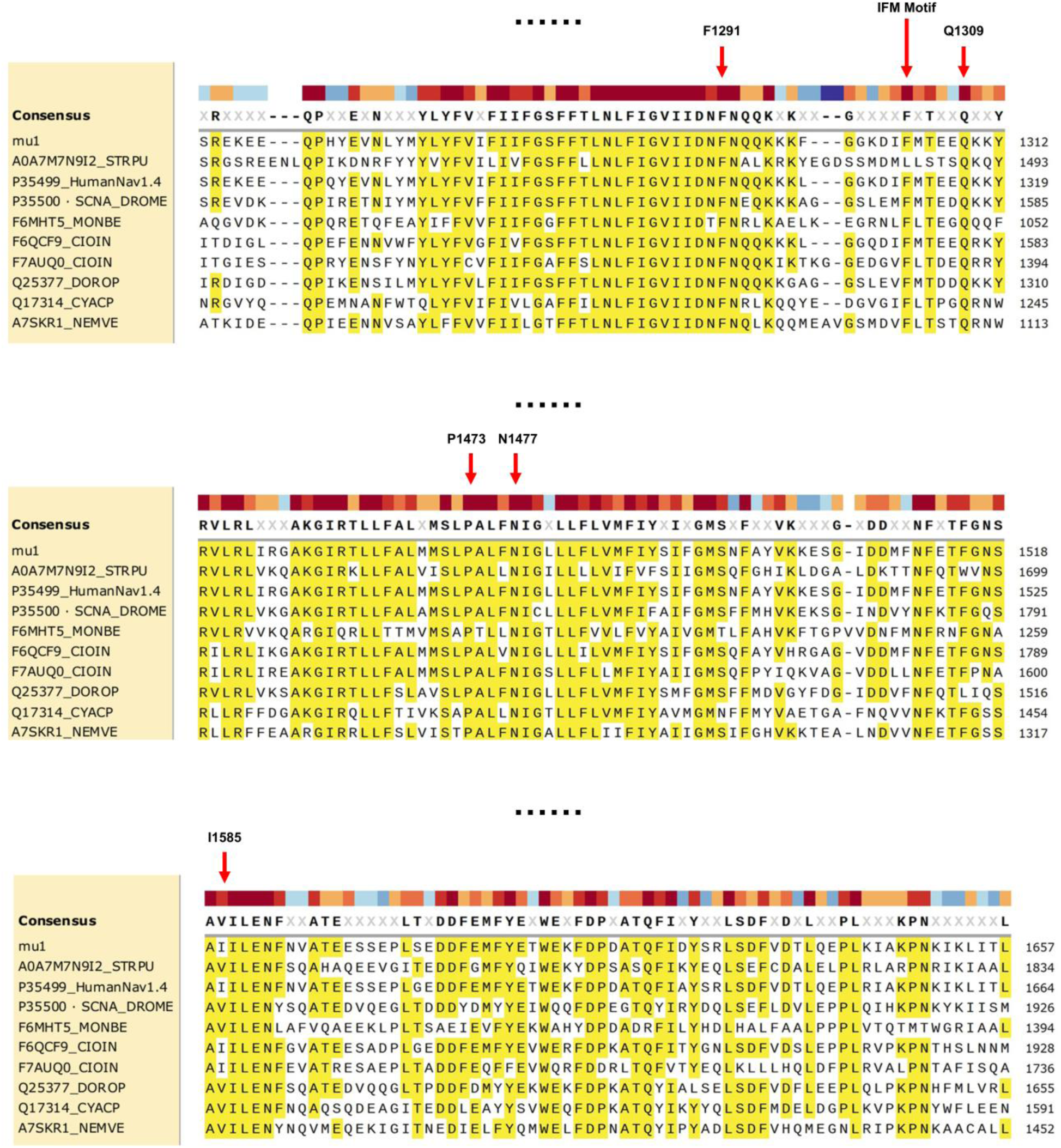
Sequence alignment of different Nav channels across the animal kingdom. Newly identified fast inactivation apparatus is highlighted by red arrows.

## Reference

1. Bean, B. P. The action potential in mammalian central neurons. Nat. Rev. Neurosci. 8, 451–465 (2007).

2. Hodgkin, A. L. & Huxley, A. F. Currents carried by sodium and potassium ions through the membrane of the giant axon of Loligo. J. Physiol. 116, 449–472 (1952).

3. Hodgkin, A. L. & Huxley, A. F. The components of membrane conductance in the giant axon of Loligo. J. Physiol. 116, 473–496 (1952).

4. De Lera Ruiz, M. & Kraus, R. L. Voltage-Gated Sodium Channels: Structure, Function, Pharmacology, and Clinical Indications. J. Med. Chem. 58, 7093–7118 (2015).

5. Fertleman, C. R. et al. SCN9A Mutations in Paroxysmal Extreme Pain Disorder: Allelic Variants Underlie Distinct Channel Defects and Phenotypes. Neuron 52, 767–774 (2006).

6. Waxman, S. G. Painful Na-channelopathies: An expanding universe. Trends Mol. Med. 19, 406–409 (2013).

7. Phillips, L. & Trivedi, J. R. Skeletal Muscle Channelopathies. Neurotherapeutics 15, 954–965 (2018).

8. George, A. L. Molecular basis of inherited epilepsy. Arch. Neurol. 61, 473–478 (2004).

9. Kyndt, F. et al. Novel SCN5A mutation leading either to isolated cardiac conduction defect or Brugada syndrome in a large French family. Circulation 104, 3081–3086 (2001).

10. Armstrong, C. M. & Bezanilla, F. Inactivation of the sodium gating current. J. Gen. Physiol. 4, 865–876 (1977).

11. West, J. W. et al. A cluster of Hydrophobic amino acid residues required for fast Na+-channel inactivation. Proc. Natl. Acad. Sci. U. S. A. 89, 10910–10914 (1992).

12. Horn, R., Ding, S. & Gruber, H. J. Immobilizing the Moving Parts of Voltage-Gated Ion Channels. J. Gen. Physiol. 116, 461–476 (2000).

13. Sheets, M. F., Kyle, J. W. & Hanck, D. A. The role of the putative inactivation lid in sodium channel gating current immobilization. J. Gen. Physiol. 115, 609–619 (2000).

14. Kellenberger, S., West, J. W., Scheuer, T. & Catterall, W. A. Molecular analysis of the putative inactivation particle in the inactivation gate of brain type IIA Na+ channels. J. Gen. Physiol. 109, 589–605 (1997).

15. Kellenberger, S., Scheuer, T. & Catterall, W. A. Movement of the Na+ channel inactivation gate during inactivation. J. Biol. Chem. 271, 30971–30979 (1996).

16. Chahine, M., Deschênes, I., Trottier, E., Chen, L. Q. & Kallen, R. G. Restoration of fast inactivation in an inactivation-defective human heart sodium channel by the cysteine modifying reagent Benzyl-MTS: Analysis of IFM-ICM mutation. Biochem. Biophys. Res. Commun. 233, 606–610 (1997).

17. Kühlbrandt, W. The Resolution Revolution. Science (80-. ). 343, 1443–1444 (2014).

18. Pan, X. et al. Structure of the human voltage-gated sodium channel Nav1.4 in complex with β1. Science *(80-. ).* **362**, (2018).

19. Huang, G. et al. High-resolution structures of human Nav1.7 reveal gating modulation through α-π helical transition of S6IV. Cell Rep. 39, (2022).

20. Jiang, D. et al. Open-state structure and pore gating mechanism of the cardiac sodium channel. Cell 184, 5151–5162.e11 (2021).

21. Jiang, D. et al. Structure of the Cardiac Sodium Channel. Cell 180, 122–134.e10 (2020).

22. Liu, Y., Bassetto, C. A. Z., Pinto, B. I. & Bezanilla, F. A mechanistic reinterpretation of fast inactivation in voltage-gated Na+ channels. Nat. Commun. 2023 141 **14**, 1–13 (2023).

23. Patton, D. E., West, J. W., Catterall, W. A. & Goldin, A. L. Amino acid residues required for fast Na+-channel inactivation: Charge neutralizations and deletions in the III-IV linker. Proc. Natl. Acad. Sci. U. S. A. 89, 10905–10909 (1992).

24. Chatterjee, A., Guo, J., Lee, H. S. & Schultz, P. G. A genetically encoded fluorescent probe in mammalian cells. J. Am. Chem. Soc. 135, 12540–12543 (2013).

25. Hyun, S. L., Guo, J., Lemke, E. A., Dimla, R. D. & Schultz, P. G. Genetic incorporation of a small, environmentally sensitive, fluorescent probe into proteins in Saccharomyces cerevisiae. J. Am. Chem. Soc. 131, 12921–12923 (2009).

26. Cha, A. & Bezanilla, F. Characterizing Voltage-Dependent Conformational Changes in the ShakerK+ Channel with Fluorescence. Neuron 19, 1127–1140 (1997).

27. Kalstrup, T. & Blunck, R. Voltage-clamp fluorometry in Xenopus oocytes using fluorescent unnatural amino acids. J. Vis. Exp. 2017, 1–9 (2017).

28. Bezanilla, F. Gating currents. J. Gen. Physiol. 150, 911–932 (2018).

29. Armstrong, C. M. & Bezanilla, F. Currents Related to Movement of the Gating Particles of the Sodium Channels. Nat. 1973 2425398 242, 459–461 (1973).

30. Chanda, B. & Bezanilla, F. Tracking voltage-dependent conformational changes in skeletal muscle sodium channel during activation. J. Gen. Physiol. 120, 629– 645 (2002).

31. Liu, Y. & Bezanilla, F. A sodium channel mutant removes fast inactivation with the inactivation particle bound. J. Gen. Physiol. 157, (2025).

32. Capes, D. L., Goldschen-Ohm, M. P., Arcisio-Miranda, M., Bezanilla, F. & Chanda, B. Domain IV voltage-sensor movement is both sufficient and rate limiting for fast inactivation in sodium channels. J. Gen. Physiol. 142, 101–112 (2013).

33. Huang, G. et al. Unwinding and spiral sliding of S4 and domain rotation of VSD. 1–9 (2022) doi:10.1073/pnas.2209164119/-/DCSupplemental.Published.

34. Galles, G. D. et al. Tuning phenylalanine fluorination to assess aromatic contributions to protein function and stability in cells. Nat. Commun. 2023 141 14, 1–12 (2023).

35. Armstrong, C. M., Bezanilla, F. & Rojas, E. Destruction of sodium conductance inactivation in squid axons perfused with pronase. J. Gen. Physiol. 62, 375–391 (1973).

36. Cole, K. S. & Moore, J. W. Potassium ion current in the squid giant axon: dynamic characteristic. Biophys. J. 1, 1–14 (1960).

37. Albert, C., Ruben, P. C., George, A. L., Fujimoto, E. & Bezanilla, F. Voltage sensors in domains III and IV, but not I and II, are immobilized by Na+ channel fast inactivation. Neuron 22, 73–87 (1999).

38. Sheets, M. F. & Hanck, D. A. Charge immobilization of the voltage sensor in domain IV is independent of sodium current inactivation. J. Physiol. 563, 83–93 (2005).

39. Berecki, G. et al. Nav1.2 channel mutations preventing fast inactivation lead to SCN2A encephalopathy. Brain 148, 212–226 (2025).

40. Armstrong, C. M. Na channel inactivation from open and closed states. Proc. Natl. Acad. Sci. U. S. A. 103, 17991–17996 (2006).

41. Horn, R., Vandenberg, C. A. & Lange, K. Statistical analysis of single sodium channels. Effects of N-bromoacetamide. Biophys. J. 45, 323–336 (1984).

42. Liebeskind, B. J., Hillis, D. M. & Zakon, H. H. Evolution of sodium channels predates the origin of nervous systems in animals. Proc. Natl. Acad. Sci. U. S. A. 108, 9154–9159 (2011).

43. Stefani, E. & Bezanilla, F. Cut-open oocyte voltage-clamp technique. Methods Enzymol. 293, 300–318 (1998).

44. Hohsaka, T., Kajihara, D., Ashizuka, Y., Murakami, H. & Sisido, M. Efficient Incorporation of Nonnatural Amino Acids with LargeAromatic Groups into Streptavidin in In Vitro Protein SynthesizingSystems. (1998) doi:10.1021/JA9813109.S001.

45. Robertson, S. A., Ellman, J. A. & Schultz, P. G. A General and Efficient Route for Chemical Aminoacylation of Transfer RNAs. J. Am. Chem. Soc. 113, 2722–2729 (1991).

46. Kalstrup, T. & Blunck, R. Dynamics of internal pore opening in KV channels probed by a fluorescent unnatural amino acid. Proc. Natl. Acad. Sci. U. S. A. 110, 8272–8277 (2013).

